# Manipulation of Seedling Traits with Pulsed Light in Closed Controlled Environments

**DOI:** 10.1101/674432

**Authors:** Shiwei Song, Paul Kusuma, Sofia D. Carvalho, Yan Li, Kevin M. Folta

## Abstract

There is substantial interest in growing crops in closed controlled environments, yet the energy requirements are high. Energy is required to produce light, but also to remove the heat generated when producing light. The goal of the current work examines a possible approach to decrease the energy requirement. The effect of pulsed light treatments was examined by monitoring seedling traits during early photomorphogenic development. Daily light integral remained unchanged between treatments, but the frequency of the pulses was varied. Developmental traits (such as inhibition of hypocotyl elongation rate) were most conspicuous during a normal photoperiod, as in twelve hours light, twelve hours darkness. Consistent with historical reports, when treatments were delivered in shorter durations (e.g. 1 hour on/off) photomorphogenic development was hindered, with the same daily light integral. However, at even shorter light intervals (e.g. seconds) seedlings developed as if they were provided full 12 h treatments. Extension of the dark period following a 5 s pulse was tested to determine the effect on seedling traits. The results showed that the dark period could be extended to at least 10 s without affecting seedling development, and extension to 20 s only had slight effects on seedling traits. The mechanism of the phenomenon was examined in Arabidopsis photosensory mutants, with substantial contributions from the phyA and cry1 pathways. The results suggest that pulsed light with extended dark periods can decrease energy input by at least 30% to >50% without affecting visible seedling traits. These pilot experiments in seedlings demonstrate that implementation of short-interval, pulsed-light strategies may lower energy requirements for growing crops in artificially illuminated environments.

## Introduction

Emerging trends in indoor farming offer alternatives to standard greenhouse or field production. Crops raised inside closed controlled environments (CCE) offer the advantage of production of high-value specialty crops in or near population centers, a low carbon footprint relative to crops shipped over distance, the use of different crop protection strategies, and opportunities to conserve water and fertilizer [1, 2]. These approaches provide profitable non-traditional production of high-value specialty crops.

However, there are challenges to growing plants in CCE, including costs of initial investment and energy use. Depending on local economic situations, energy alone represents between 25% [3] and 60% [4] of production costs, and they have a greater carbon footprint than traditional greenhouses [5]. These realities present bottlenecks to profitable production, as energy is needed for lighting as well as to dissipate the heat energy generated from light sources [reviewed in 6].

Light is required for plant growth via photosynthesis, but it also has an important role in shaping plant physiology, development, metabolism and morphology during the developmental program called photomorphogenesis [reviewed in 7, 8]. Because plants are rooted in one place, plants must continually monitor the ambient environment and adjust gene expression patterns to best acclimate to prevailing conditions [9, 10]. The light environment is monitored through a series of specialized light sensors, each with an optimal range of activation, and each with downstream signaling pathways that terminate in mechanisms that shape plant biology. Over the last decade many reports have examined the effect of narrow-bandwidth spectrum, photoperiod and fluence rate with the intent of manipulating plant traits in controlled environments [11]. However, while control of light quantity and quality has become exquisite, the energy needs to drive plant production remain high.

Clearly light quantity (fluence rate), quality (wavelength), duration (photoperiod) can be altered individually or in combination to influence plant traits using today’s sophisticated technologies. In this report we revisit another variable–the frequency of pulses applied in intervals of seconds to hours. While mostly confined to studies of intermittent pulse rate and photosynthesis [12, 13], other historical reports noted that varying treatments from seconds to minutes to hours can have profound effects on plant growth and development, with seconds-long pulses being visibly indistinguishable from a normal 12 hour dark-light photoperiod [14].

Because plants grow well in seconds-long pulses, it becomes possible to explore energy savings by extending the dark period between pulses. The hypothesis is that a defined frequency of pulses may lead to production of identical plant products with fewer photons invested. These experiments exploit two factors not available to Garner and Allard in 1931-- the known plasticity of highly-responsive seedlings, photomorphogenic mutants, and narrow-bandwidth light sources. Here these three tools have been combined to examine the effects of light pulses on growth and development, with special attention to the length of the dark period between pulses. Consistent with eighty-eight-year-old observations using full-spectrum white light [14], the results indicate that seedlings treated with pulse light for several seconds are indistinguishable from those treated with a normal photoperiod. Examination of the dark interval shows that it may be extended without affecting seedling development for well past the equivalent seconds-long treatment to obtain the same effect from one 12 h light treatment. The use of pulsed light with extended dark periods provides an additional tool to manipulate plant growth and development, and may provide energy savings by supporting comparable growth with significantly lower daily light integrals.

## Materials and Methods

### Plant Materials

Red Russian Kale, Purple Top Turnip, and Ruby Queen Beet Seeds were purchased from Johnny’s Selected Seeds (Fairfield, ME). Arabidopsis experiments were performed with seedlings in the Columbia (Col-0) background. Experiments for cryptochrome (*cry1-304* and *cry2-1*) mutants were performed in the Col-0 background. Phytochrome mutants (*phyA-201* and *phyB-5*) were tested in the Lansberg *erecta* (Ler) background.

### Growth and assay conditions

Survey experiments were performed on seeds placed on autoclaved BioStrate Felt saturated with 0.1x Hoaglands basal salt hydroponic solution inside glass, wide-mouth, pint canning jars covered with the lower half of a plastic petri dish. The seeds were stratified in darkness at 4°C for 72 h, and then transferred to room temperature (22°C ±2°) and exposed to cool white fluorescent light (15 μmol m^−2^ s^−1^) for 1 h, and then moved into complete darkness for a minimum of 2 h before being transferred to the specific light treatment.

Pulse-light experiments were performed on seedlings on an agar substrate in plastic Magenta boxes. Twenty-five Red Russian Kale (*Brassica napus*), Purple Top Turnip (*Brassica rapa*, Mountain Valley Seed Co.), or Ruby Queen beet (*Beta vulgaris*, Eden Brothers) seeds were surface sterilized with a rinse in 70% ethanol in a laminar flow hood, and then placed in a grid pattern on ½ x MS medium (pH5.8) solidified with 0.75%-0.80% phytoagar. Seeds were then stratified for 72 h, given a 1 h white light treatment (15 μmol m^−2^ s^−1^) to synchronize germination, and then were moved to specific light conditions. Arabidopsis experiments were performed in identical conditions except that seedlings were grown on square Petri dishes oriented vertically, allowing seedlings to grow up the surface of the plates for ease of imaging.

Narrow-bandwidth LED light treatments were provided by Plant Whisperer MarkV light units (Light Emitting Computers, Victoria, BC Canada). Wavelengths tested include 450 nm (blue; B), 660 nm (red; R), and 730nm (far-red; FR). For some light pulse experiments both a GRALab model 655 intervalomter and the Apollo 6 Timer (Titan Controls) were used. In later experiments pulse-light treatments were performed using the custom LED light sources where the DC limb was controlled using a custom Raspberry Pi based electronic relay array. Trials were conducted in ventilated growth chambers lined with reflective mylar maintained at 22°C (±2°C). Light treatments varied in wavelength, pulse duration, total hours of light exposure, and fluence rate as described below.

After 96 h of treatment time, fresh weight was measured before the seedlings were transferred to a flat-bed scanner for image capture for hypocotyl length measurement. The seedlings were then frozen with liquid nitrogen, and crushed into a powder using a mortar and pestle. The powder was then placed into a 1.7mL microcentrifuge tube for anthocyanin measurements. Roughly 100mg was used for each extraction.

For anthocyanin measurements 300 μL of methanol-1%HCl was added to each tube, and then placed in darkness at 4°C for at least 16 h [15]. Next, 200 μL of water and 500 μL of chloroform were added to the tube, the contents were vortexed and then centrifuged in a microcentrifuge at the highest speed for 5 min. Roughly 400 μL of the supernatant was moved to a new tube, to which 400 μL of 60% methanol-1%HCl 40% water was added. A SmartSpec 3000 spectrophotometer (Bio-Rad) was used to measure the absorbance at 530 and 657 nm. 60% methanol-1%HCl 40% water was used as a blank. Final anthocyanin per mg tissue was calculated using the following equation: Anthocyanin/mg FW tissue = ((Abs_530_−Abs_657_)×1000)/powder weight (mg)

### Statistical analysis

All experiments were repeated at least three times. Statistical analyses were performed using the statistical software package SPSS 19.0 (SPSS, Inc., Chicago, IL, USA). Data were analyzed by one-way analysis of variance with LSD method.

## Results

### Developmental Response to Light Treatment

To examine the effect of different light pulse lengths at a common daily light integral (DLI), seedlings from three genotypes were placed into chambers and treated with various fluence rates of far-red light. FR was chosen because it induces strong responses in Red Russian kale and is widely recognized for its strong effect on inhibition of hypocotyl elongation and anthocyanin accumulation [16]. At the same time FR light sources do not significantly excite photosynthesis, so using these wavebands allows uncoupling of developmental response from influences arising from chloroplast development and photosynthesis. Seedlings were grown for 96 h under pulses of FR light, varying from 5 s to 12 h, all with a fluence rate 100 μmol m^−2^ s^−1^ and DLI of 4.32 mol m^−2^. The results are presented in in Figure 1.

**Figure 1.**
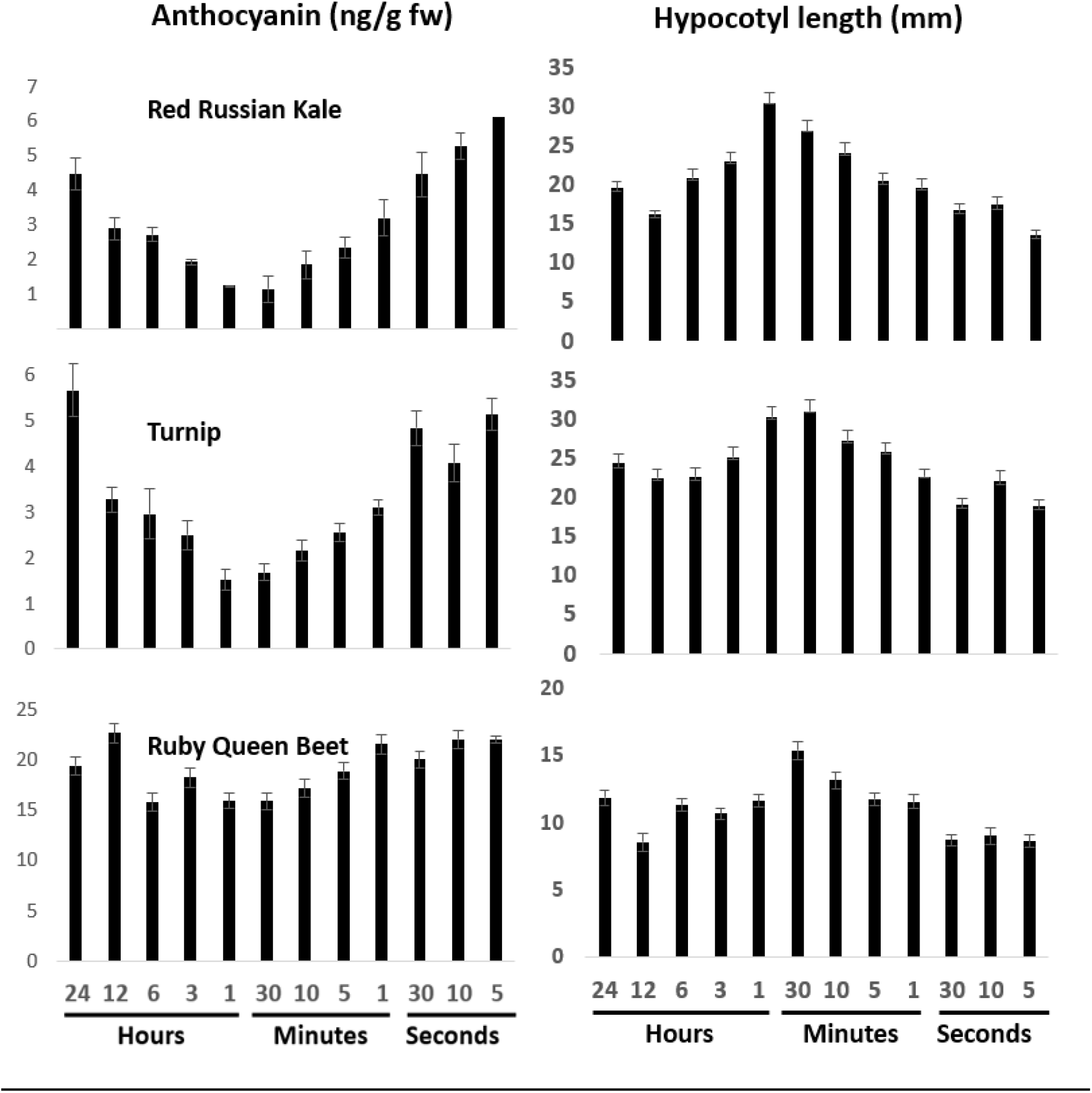
Light pulse frequency effects in select seedlings. Inhibition of hypocotyl elongation and anthocyanin accumulation were assessed in seedlings of Red Russian Kale, Purple Top Turnip, and Ruby Queen Beet in response to 10 μmol m^−2^ s^−1^ far red light, applied for various time intervals for 96 h. The mean represents measurements from one experiment with at least 30 seedlings. Independent experiments were performed with similar results. Error bars represent standard error of the mean.

The results show that in Red Russian Kale and Purple Top Turnip seedlings accumulate anthocyanins in a 12 h (on/off) light interval. As the light interval is split into smaller time periods there is less accumulation as pulse times decrease from 6 h down to 1 min, with the lowest accumulation occurring between 30 min and 1 h. Light pulses of 30 s and under led to the same amount of anythocyanin (or possibly more) as a 12 h treatment. The Ruby Queen beet also exhibited a similar trend, but the absolute levels of anthocyanin were higher than in the other two seedlings. Further experiments would focus only on kale and the turnip seedlings.

A similar trend was observed for hypocotyl elongation (Figure 1). Seedlings grown under longer intervals that approximate normal day/night intervals exhibited the shortest hypocotyls, with the longest hypocotyls typically occurring in response to pulses from 10 min to 1 h. However, shorter pulses (5, 10, 30 s) produce the same outcomes as the longer 12-h pulse.

Fresh weight, chlorophyll accumulation and anthocyanin accumulation were all measured under these conditions. All of these measurements showed trends in response to light that were highly similar to those observed for hypocotyl length (K. Folta, P. Kusuma; Unpub. obs.). These observations indicated that hypocotyl elongation was a suitable proxy for measuring photomorphogenic progression. Further experiments would focus strictly on hypocotyl growth inhibition under various light treatments.

### Persistence of Light Response

The data from Figure 1 show that seedlings receiving the same DLI spread over 12 h or 5s intervals are comparable in terms of development and physical attributes. It was therefore of interest to examine the persistence of the light effect after the light pulse ended, and determine how long the dark period may be extended before the photomorphogenic development slowed. At the same time, growth under different wavebands may inform how different photosensory systems are contributing to photomorphogenic development along with their kinetics of dark reversion. To test these attributes, kale and turnip seedlings were grown under 5 s pulses of R, B or FR light (~140 μmol m^−2^ s^−1^ PAR; B= 30 μmol m^−2^ s^−1^, R= 110 μmol m^−2^ s^−1^, FR= 30 μmol m^−2^ s^−1^) and the dark period was extended. The results of at least three independent experimental replicates are presented in Figure 2.

**Figure 2.**
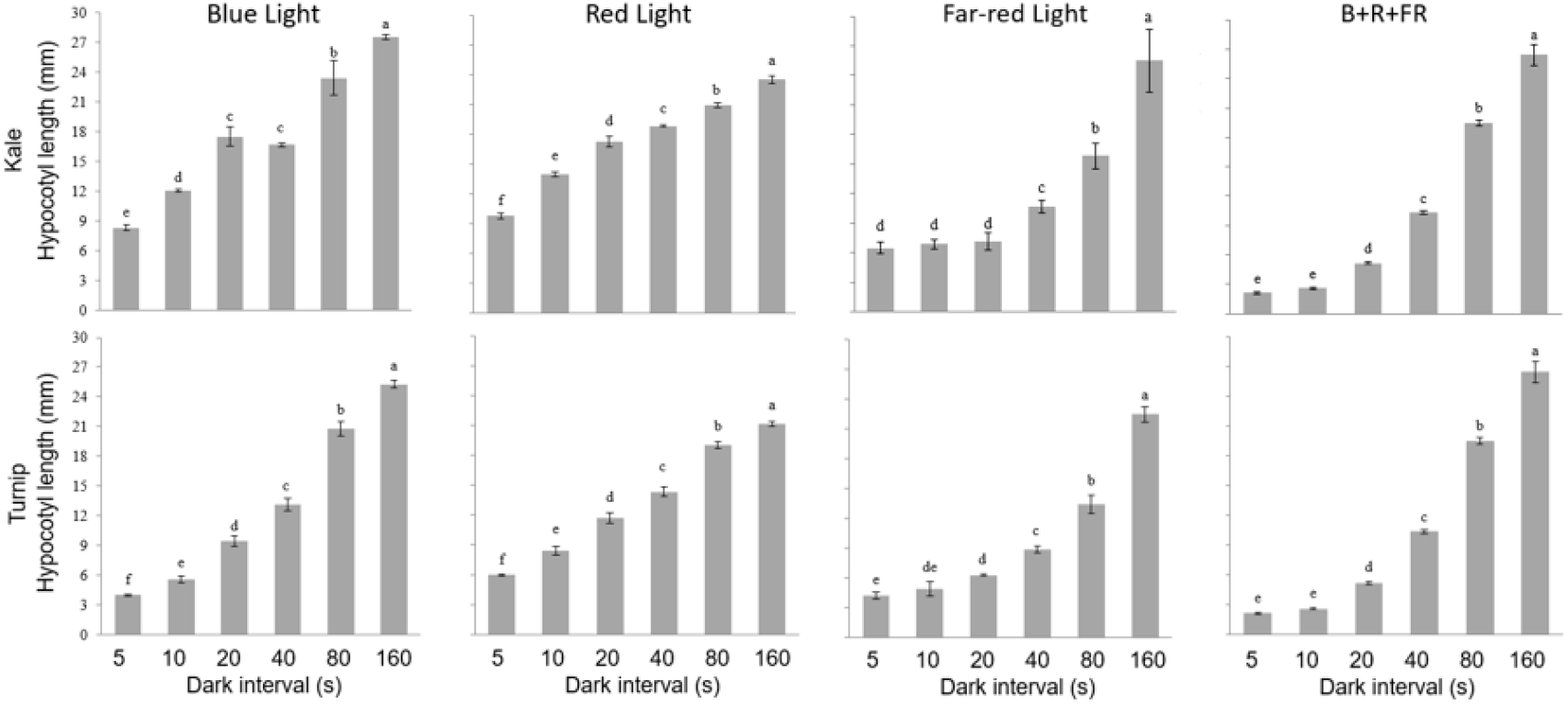
The effect of extending the dark period following a light pulse in Red Russian Kale and Purple Top Turnip. Inhibition of hypocotyl elongation was assessed in response to 100 μmol m^−2^ s^−1^ blue, red, far-red, or combination (B:R:FR= 1:4:1; ~140 μmol m^−2^ s^−1^) light, followed by a dark period ranging from 5 to 160 s for 96 h. For each experiment N>20 seedlings and the mean reflects the results of three independent experiments. Error bars represent the standard error of the mean. Different letters represent significant differences by LSD ANOVA analysis (p<0.05).

B and R light effects on hypocotyl growth inhibition begin to revert 10 s after a 5 s pulse in both kale and turnip. The effect of FR illumination was more persistent, as growth inhibition did not revert significantly until after 20 s of darkness in kale, and small (yet statistically significant) differences were seen in turnip seedlings after a 10 s dark period. A combination of light inputs (B= 30 μmol m^−2^ s^−1^, R= 110 μmol m^−2^ s^−1^, FR= 30 μmol m^−2^ s^−1^) provided stronger inhibition that persisted until after 10 s, but was still comparable to single-waveband 5 s dark treatments after 20 s of darkness. Trends of anthocyanin and chlorophyll accumulation were similar, indicating that the plants were not significantly compromised by the treatment, but were accumulating fewer pigments in response to light.

### Experiments in Arabidopsis

Seedlings from the model plant *Arabidopsis thaliana* were examined for their response to the same pulse conditions in Figure 3. Repeating the experiments in Arabidopsis allows examination of the response in a well-characterized physiological system and opens the possibility to examine mutant genotypes that may aid in understanding the genetics of persistent light signaling when plants are treated with extended dark periods.

**Figure 3.**
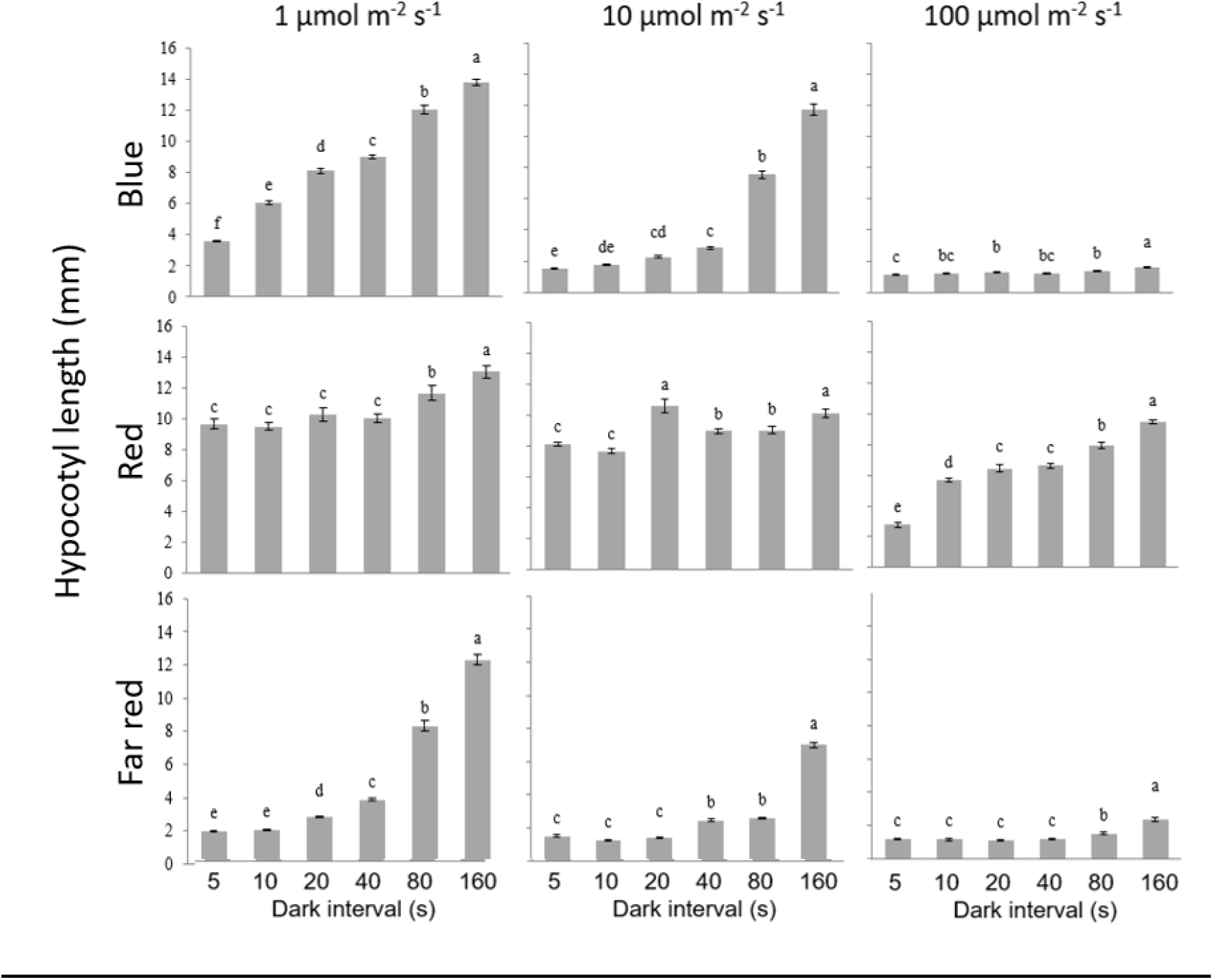
Wavelength dependence and fluence-rate/response effect on hypocotyl inhibition. *Arabidopsis thaliana* seedlings were germinated and grown under various pulse conditions under various fluence rates of blue, red or far-red light. Light pulses were applied for 5 s and the dark interval varied. Hypocotyl lengths were assessed after 96 h. The means represent outcomes from three independent experiments (N>20 each). Error bars represent the standard error of the mean. Different letters represent significant differences by LSD ANOVA analysis (p<0.05).

The results in Figure 3 show the Arabidopsis (Col-0) response to pulse light with increasing dark periods after different fluence rates of B, R, and FR light. Seedlings treated with 5 s pulses at 1 μmol m^−2^ s^−1^ B light exhibit strong growth inhibition, but the response reverts quickly, as seedlings show loss of inhibition after 10 s of darkness, and a relatively linear loss with increasing dark intervals. Seedlings treated with 10 μmol m^−2^ s^−1^ retain most of their inhibition even after 40 s of darkness. Illumination with 5 s of B light at 100 μmol m^−2^ s^−1^ leads to strong growth inhibition that is almost completely retained to 160 s.

To the contrary, R light pulses had a much lower effect on growth inhibition in amplitude, yet the effects were persistent even at low fluence rates. The effects of FR light pulses were much like those of B light, with a strong effect on growth inhibition that was maintained after 20 s of darkness, even at low fluence rates. At 100 μmol m^−2^ s^−1^ the inhibition was persistent, unchanged after 40 s of darkness, and maintaining the majority of growth inhibition even up to 160 s of dark period.

### Genetic Analysis

The experiments outlined in Figure 3 indicate that different photosensory systems vary in maintaining the response to the pulse treatment. The mechanism can be better explored using photomorphogenic mutants in Arabidopsis. The experiments in Figure 3 were repeated using co-irradiation with R, B and FR light, tested on the *phyA*, *phyB*, *cry1*, and *cry2* mutants and their corresponding wild-type genotypes. Three wavebands were delivered simultaneously (as described above) to observe the effects of coaction. The results are shown in Figure 4.

**Figure 4.**
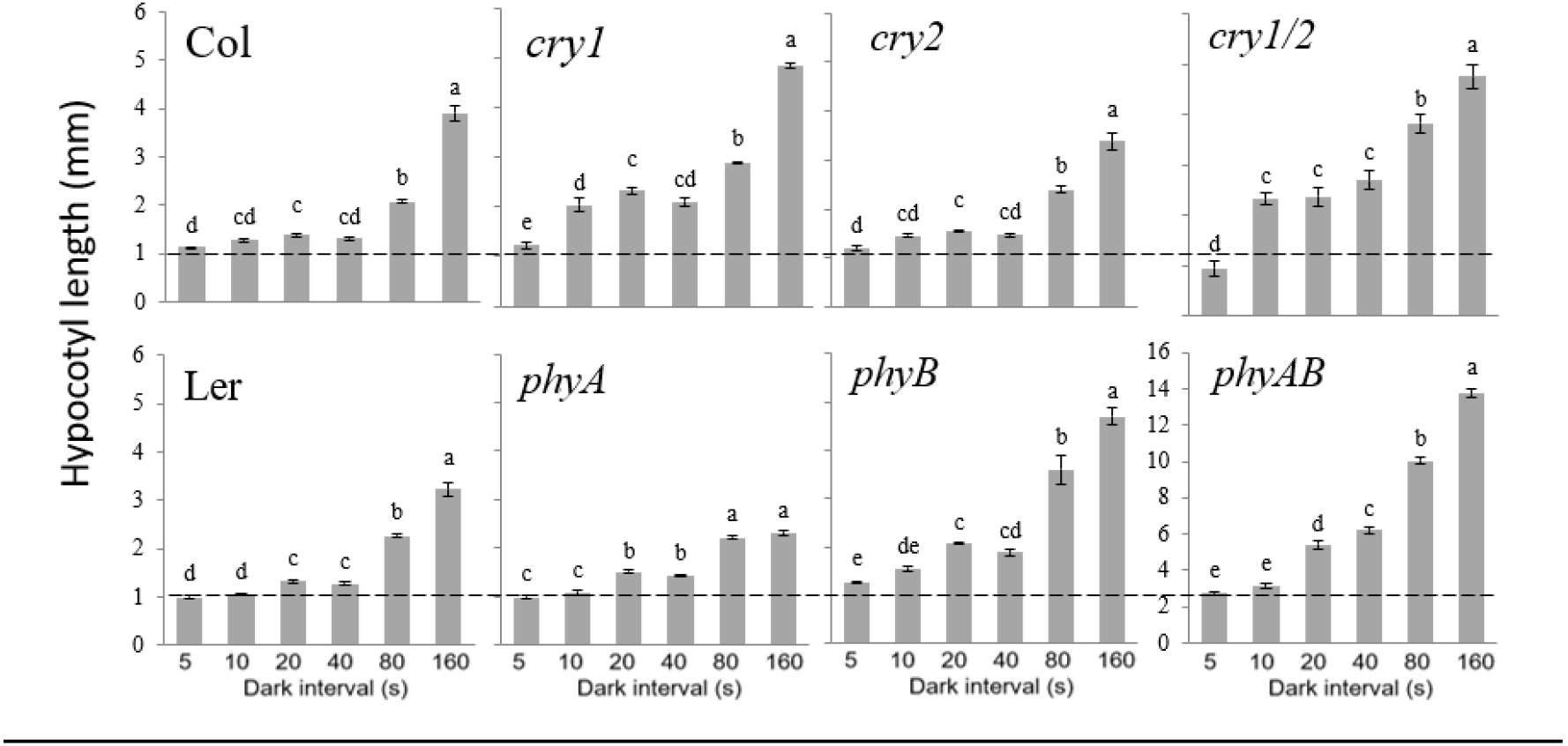
The effect of light pulse persistence in Arabidopsis mutants and corresponding wild-type plants. Arabidopsis seedlings were grown under a combination of blue, red, and far-red light (1:4:1) and a fluence rate of ~140 μmol m^−2^ s^−1^, pulsed for five seconds with a variable dark period. The data represent averages of three independent experiments (N>20). Error bars indicate standard error of the mean, and letter designations represent significant differences by LSD ANOVA analysis (p<0.05).

The first observation is that both Col-0 and Ler genotypes respond strongly to all three wavebands in combination, reaching what appears to be full inhibition and retaining hypocotyl growth inhibition at least until 40 s of darkness. Examination of photomorphogenic mutants shows that cry2 and phyA receptors are not required for retention of maximal growth inhibition, and that the absence of cry1 or phyB leads to lower levels of inhibition in the 5-40 s dark interval treatment. However, the loss of either receptor does not lead to complete loss of response under illumination with multiple bandwidths. Examination of the double *cry1cry2* and *phyAphyB* mutants shows that the effects of double mutants are approximately the same as loss of either the cry1 or phyB receptor alone. At longer dark intervals (80 or 160 s) *phyA* mutants still show inhibition that is not observed in *phyB* or *phyAB* mutants.

## Discussion

The realistic potential of closed controlled environment (CCE) agriculture is limited by several factors, which include high costs of energy for illumination and temperature management [17]. The adoption of these technologies is tightly tied to grower profit, and that is substantially dependent on minimizing electricity costs. One of the other frequent justifications of the technology is that there is a potential to eliminate the carbon footprint of shipping to major urban centers, yet that is a claim that has yet to be realized, as CCE operations require substantial energy input [3, 5, 18].

Several strategies have been proposed to decrease the energy demands of CCE. Plant breeding efforts are developing new varieties that perform better in low-light environments [19]. Robotic mobile lighting systems produce intervals of illumination close to the plants, and produce the same amount of butterhead lettuce while cutting energy requirements in half [20]. Other groups have cut energy costs using lensing on LED light fixtures to focus radiant energy on leaves rather than between plants [17]. The work outlined in this report explores the use of pulsed illumination, from seconds to hours, to shape informative seedling traits. The dark period was then extended to examine how a light signals persist in darkness, lowering DLI and energy input.

The experiments follow observations from Garner and Allard [14] where they examined the effects of abnormal photoperiod on plant performance. In their report plants were grown under high intensity white light, with light/dark intervals ranging from 6 h down to seconds. The authors observed that plants treated with intervals from 1 h to 1 min performed poorly and exhibited symptoms of malnutrition, such as less chlorophyll and less accumulation of dry matter. However, plants grown under 5 s intervals performed the same as normal 12 h photoperiod controls [14]. In all of their conditions the pulses induced flowering in long-day plants and inhibited flowering in short day plants, indicating that the treatments were sufficient to satisfy a daylength requirement.

In the present study the principles of this seminal work were applied to developing seedlings. The work used seedlings because the developing seedling is exquisitely sensitive to light signals. Light sensing, integration and response results in conspicuous morphological, molecular and biochemical changes that may be easily monitored and measured. Traits such as inhibition of hypocotyl elongation, light mediated gene expression and the accumulation of photosynthetic or screening pigments are closely tied to light input, and respond proportionately over a large range of fluence rates and wavelengths. The experiments may be performed over the course of 96 h, allowing examination of a large number of seedlings in a short time. Principles identified in seedlings may later be tested in mature crop plants.

The species tested offer specific strengths and limitations. Red Russian Kale and Purple Top Turnip have been previously characterized and possess several conspicuous seedling responses to light across the spectrum, with a long range of linear response to varying fluence rates (P. Kusuma, K. Folta, unpub. obs.). While performing well in these experiments, the selection of two highly-plastic brassicas frames a central limitation to interpreting the results of the trials, as it is unclear how such treatments translate to distant taxa. The findings did translate well to two *Arabidopsis thaliana* ecotypes. While still brassicas, they are not crop plants, so it suggests that the ability to integrate an intermittent light signal is not something that arose from selection and is likely inherent to seedlings, at least within brassicas. Future experiments will test these responses in other important seedling varieties used as commercial food crops as well as in mature plants.

The basic experiments show that seedlings grown under 5 s on/off pulse are roughly comparable developmentally to seedlings grown under 12 h on/off pulses (Figure 1). The DLI is exactly the same at a given fluence rate (4.32 mol m^−2^). Intervals that are 30 min or 1 h min long show less photomorphogenic advancement, despite treatment with the same DLI. As treatment durations increase to 3 and 6 h, plants show more photomorphogenic progression, before reaching their expected phenotype at 12 h on/off. Under all conditions the differences in photomorphogenic development reflect effects of the timing of light/dark cycles. These findings begin to touch on the mechanism for how pulsed treatments work, as these results indicate that seedlings must have a means to gate light input into developmental processes that limits signals when they are delivered in pulses that are 30 min to 1 h in duration.

A good candidate might be components that input into the circadian oscillator. For instance, the Evening Complex [21] is a multi-protein complex formed from interaction of EARLY FLOWERING3 [22] EARLYFLOWERING4 and LUXARRYTHMO [23]. It is expressed rhythmically and interacts with phyB. Ultimately the complex connects the internal oscillator to processes such as expression of genes associated with photosynthesis and plant growth. Developing seedlings with defects in the Evening Complex exhibit longer hypocotyls [23, 24]. The hypothesis is also consistent with the concept of “gating”, a process where a response cannot be activated because a molecular-biochemical constraint restricts the magnitude of circadian-oscillator conditioned responses [25].

It is tempting to speculate that short pulses are insufficient to condition the oscillator (override the gate), so from a photosensory viewpoint seedlings interpret short (seconds) pulses as constant light. However, when pulses reach 30 min to several hours in duration it is sufficient to set the biochemical complexities of the internal oscillator in motion, only to be disturbed by an unexpected period of darkness. The chloroplast also may be disturbed, as photosynthetic rates and partitioning of resources now occurs in intervals that are not in concert with the internal oscillator.

Because the 5 s on/off treatments are inconsequential to seedling growth, it was of interest to extend the dark period to determine how long the light could be removed without affecting seedling traits. The extended dark period could cut DLI, yet potentially yield the same seedling outcome. Seedlings were then grown under light-dark intervals where the dark period was extended to 10, 20, 40, 80 and 160 s. The results of the treatment illustrate the decay of the photomorphogenic response with time in Red Russian Kale and Purple Top Turnip (Figure 2). Far-red light exerts effects that are interval dependent and demonstrates the persistent activation of far-red light signaling after treatment. Far-red light has been shown to exert powerful effects on inhibition of stem elongation [26, 27] and also promotes anthocyanin accumulation [28–30], and therefore is a way to uncouple developmental cues from photosynthetic influence. The effects of pulses on these processes confirms that at least part of the effect being observed is due to excitation of photomorphogenic systems, namely phyA. The cry1 receptor has been shown to be active for at least six minutes [31]. The persistence of a response to a 5 s pulse after 80 s darkness demonstrates that all relevant photoreceptors are saturated and excess energy is not able to contribute to the response. In photobiological terms reciprocity failure observed, as there is a non-linear response between photons applied and response measured. The likely cause is the molecular or physiological bottleneck that occurs when the seedling cannot sense, signal, or respond any further to an increasing level of input. This is an important threshold to understand, as energy applied once the plants have achieved a full response is essentially wasted.

This information is important in the design of artificial lighting environments. At least in seedlings under B, R and FR treatment, the coaction of multiple sensory systems leads to a more complete suppression of hypocotyl elongation than any single treatment alone. Figure 2 shows that the combination of B, R and FR leads to inhibition that is stronger than any single waveband, although the influence of the signal wanes after 10 s of darkness, and seedlings were substantially longer with 20 s of darkness following the light treatment. Supplemental Figure 1 shows the effects on pigments, as anthocyanin and chlorophyll levels slow diminish with the extended dark period, yet not to the degree that hypocotyl elongation rate is affected. It also is important to note that both Red Russian Kale and Purple Top Turnip were likely selected because of their purple color, so anthocyanin levels may be expected to be high. There may be much less in other seedling varieties, but the results show that many traits may still be maintained through significant periods of darkness. Such findings could have a profound effect on energy savings if implemented in production contexts.

The same responses were tested in the model plant Arabidopsis thaliana. These tests were appropriate because of the substantial photophysiological and genetic understanding of inhibition of stem elongation in the species. The results of narrow-bandwidth light treatments indicate that a relatively low fluence rate treatment of 10 μmol m^−2^ s^−1^ B or FR light can strongly inhibit stem elongation, and that the response persists even after 40 and 80 s respectively. Treatment with 100 μmol m^−2^ s^−1^ leads to potent suppression that persists even after 160 s for both wavelengths. The data indicate that the treatments approximate a dose-response relationship over a limited number of points tested. More importantly, a 5 s pulse of FR light at 1 μmol m^−2^ s^−1^, applied every 15 s is sufficient to suppress stem elongation almost maximally. These findings introduce the basis of examining the photon economy of plant development, as a relatively low fluence rate treatment more frequently may be more effective than a less-frequent high-fluence-rate pulse.

The results also show that R light sensing systems are less responsive than FR or B. Strong effects of R light on hypocotyl elongation rate are only seen with frequent pulses of high fluence rate light. At least with respect to Arabidopsis seedlings, and the limited data from kale and turnips, these results suggest that R light may be dispensable in maximizing early developmental response to light. B and FR light are more potent per photon delivered, which is an important consideration in the design of lighting regimes for plant growth. Light sources for these wavebands have approximately the same output in terms of photons per joule (Pattison et al., 2018), with substantial differences in how that energy investment translates to plant development. An energy honing strategy may be to provide a 1 μmol m^−2^ s^−1^ pulse of FR light every 10 s, and a 10 μmol m^−2^ s^−1^ of B light every 40 s. Robust suppression of stem elongation rate is achieved with fewer photons invested that is similar to a single daily high-fluence-rate 12 h treatment.

The results are consistent with what is observed in genetic analyses, which were performed on seedlings grown under blue, red and far-red light in combination. The results in Figure 4 show that elimination of cry1 leads to reversion of the sustained inhibition, and that cry2 has little influence. With respect to phytochromes, phys appear to be acting redundantly, and even the *phyAphyB* mutant shows inhibition after 10 s of darkness, likely an effect imparted by *cry1*. It is also worth noting that elimination of phyA leads to stronger growth inhibition after 80 s of darkness, suggesting a role in growth promotion that is opposing the effect of phyB, and other receptors.

The next steps will attempt to translate these results in seedlings to mature plants. Tests will examine if pulsed-light treatments can drive production of microgreens or even leafy greens with results comparable to normal-photoperiod plants. Even if the principles shown here fail to translate to mature crops the use of light interval treatments clearly is effective in seedlings, and the capacity to conserve energy during establishment is still an important gain even if the plants require normal photoperiodic treatments going forward. It also may be possible to select specific varieties that perform well under these light-dark treatments with extended dark periods. In all cases these seconds-long pulses may be one other way to limit the cost of production in closed, controlled environments.

## Acknowledgements

The authors thank the Chinese Scholarship Council for supporting the visiting scientist appointment of SS. SC’s postdoctoral appointment was funded by the United States Department of Agriculture Grant number. We thank by Light Emitting Computers (Victoria, BC Canada) for providing summer salary for PK. We present this work in memory of LEC founder Jeffery Bucove who shared a vision of using artificial lighting strategies to improve plant production to aid the human condition.

## Supplemental Figure

**Supplemental Figure 1.**
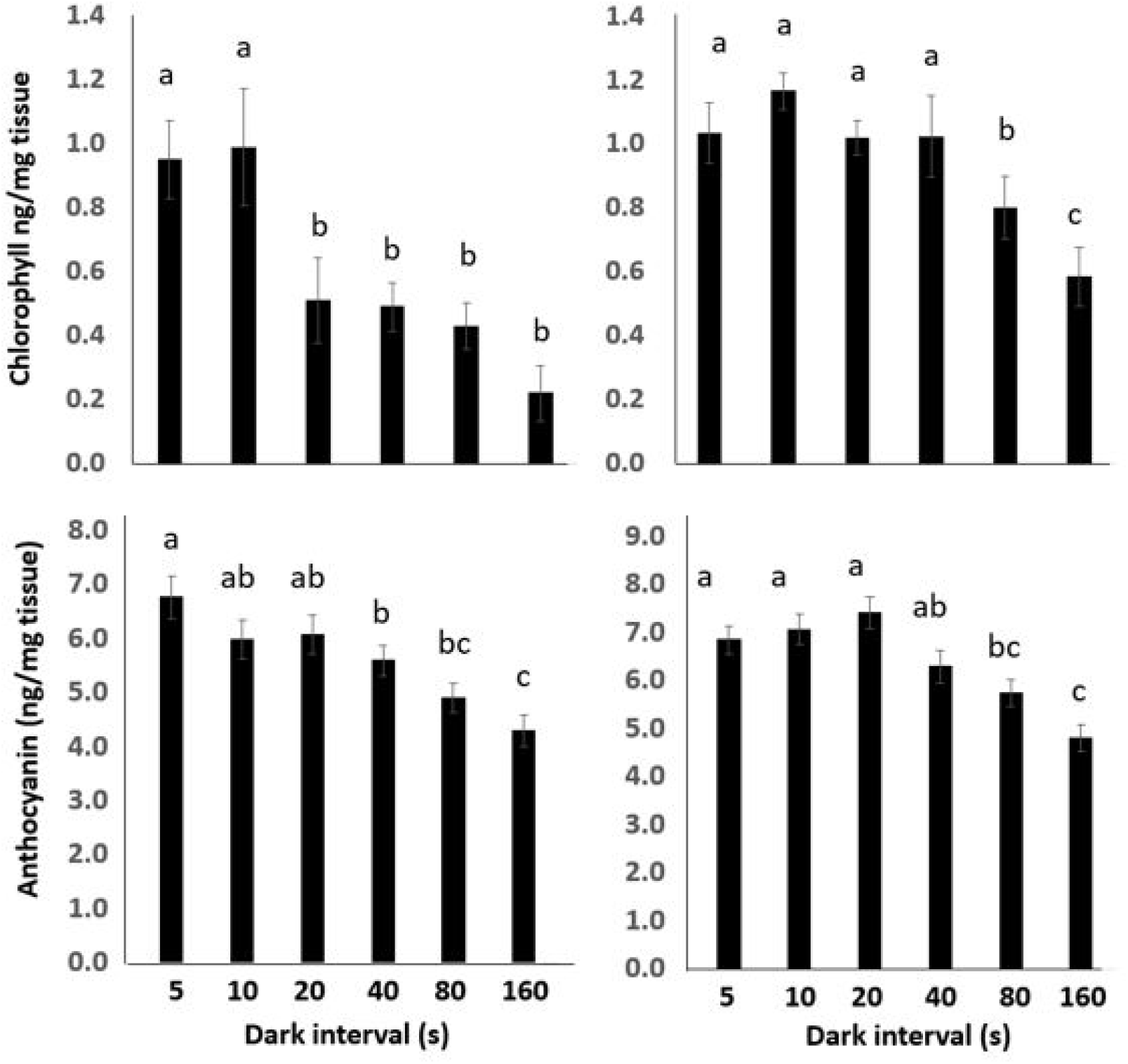
The effect of extending the dark period on pigment accumulation in Red Russian Kale and Purple Top Turnip. Anthocyanin and chlorophyll accumulation levels were anlayzed in response to 100 μmol m^−2^ s^−1^ blue, red, far-red, or combination (B:R:FR= 1:4:1; ~140 μmol m^−2^ s^−1^) light, followed by a dark period ranging from 5 to 160 s for 96 h. For each experiment N>6 seedlings and the mean reflects the results of three independent experiments. Error bars represent the standard error of the mean. Different letters represent significant differences by one-way ANOVA with Tukey HSD post-hoc test (p<0.05).

